# Evaluation of radioprotective effect of chickpea (*Cicer arietinum*) in the survival of *Lasioderma serricorne* (Fabricius, 1792) (*Coleoptera*, *Anobìidae*) irradiated with ^60^Co

**DOI:** 10.1101/637322

**Authors:** Adilson C. Barros, Kayo Okazaki, Valter Arthur

## Abstract

We investigated the presence of natural radioprotectors in food using a new technical modality that utilizes the insect *Lasioderma serricorne* as a radiosensitivity bioindicator to check radioprotection properties in minimally processed chickpeas. The insects were obtained from the entomological biotherium of the Laboratory of Radiobiology and Environment of CENA-USP. They were fed with an experimental diet and just when the first generation hatched completely, the experiments were conducted. The randomly chosen control diet, consisted of three parts of wheat germ, one part of brewer’s yeast, and a slice of French bread toasted in an oven previously set up for humidity control. The diet of chickpeas consists only of whole grains crushed in a mechanical grinder to obtain flour. The result was significant for the survival of insects (p<0.0001) reared on a diet of chickpeas compared to those reared on control diet irradiated with gamma rays from ^60^Co in the range of 5.0 to 1500 Gy. We presented statistical evidence that the chickpea diet has radioprotective properties in the insect for gamma rays.

**SUMMARY STATEMENT:** The study is important because it shows that chickpea has protective properties against ionizing radiation, how to act against its biological effects and minimize them.

## 1. INTRODUCTION

The radiological protection of an organism can be physical or chemical. Physical protection against ionizing radiation occurs through time, distance, shielding and contamination control (Bittelli, 2006). From the chemical point of view, the agents that are able to minimize the deleterious effects of ionizing radiation in cells or organisms are known as radioprotectors. The radioprotectors may be synthetic or natural and are able to minimize the deleterious effects of ionizing radiation in an organism, if previously absorbed by organs and tissues in evidence. The present study arose from a casual observation in the laboratory of Radiobiology and Environment in the CENA-USP, Brazil, whose goal is the disinfestation of pests. It was observed that the same pest showed different survival responses when fed with different types of grains under the same condition of disinfestation by radiation (type of radiation, dose and dose rate) (Arthur et al., 2012). In addition, there are reports about Japanese survivors of the atomic bombs, whose rate of incidence of leukemia and the latency period varied depending on alimentary habits of each community. There is a need to protect humans against the harmful effects of ionizing radiation (Jagetia, 2007a). The use of chemicals to protect against the harmful effects of radiation was attempted after World War II with the goal to safeguard humans against the military use of atomic weapons. Patt and his co-workers (1949) were the first to investigate the effect of amino-acid cysteine in rats exposed to lethal doses of X-rays. The radioprotectors currently used are all synthetic and cause collateral effects in radiotherapy applications. A natural radioprotector should be easily available. Dietary agents that are already consumed by humans have not received the attention they deserve for their potential radioprotective effects (Jagetia, 2007b). Certain chemicals are able to protect against the harmful effects of radiation, but more than 50 years of research has produced only one radioprotective drug approved by FDA: WR-2721 or amifostine. Amifostine is a synthetic aminothial radioprotector used in radiotherapy that has some side effects such as toxicity, sneezing and drowsiness, but it does not protect the brain because the drug cannot cross the blood-brain barrier and also does not show to strongly protect the lung. (Hall, 2000). This drug offers good protection, but is relatively toxic (nausea, vomiting and hypotension being some of the most common adverse effects) (Hosseinimehr, 2007). The general utility of amifostine is limited by its inherent toxicity and high cost (Devi and Agrawala, 2011). This paper proposes to identify radioprotective principles of natural occurrence, namely in food, whose side effects are minimal or nonexistent and which are easily available. The importance of this work is due to the possibility of exploring new methods of radiation protection at the occupational and therapeutic level, as well as to establish a new methodology for this research category. The insect *Lasioderma serricorne* (*Coleoptera*, *Anobiidae*) was used as the bioindicator of radiosensitivity due to its wide availability, easy handling, low cost, compact space compared to *rodentia* models, and availability of a large number of individuals for the experimental plan. A severe DNA fragmentation in *Lasioderma serricorne* cells, whose adult form shows higher resistance to radiation, was observed just after irradiation, with the damage being repaired during the post-irradiation period in a time-dependent manner (Kameya et al., 2012).

The insects of Anobìidae family are cylindrical or oval beetles, 1-9 mm long and have a head which is bent down under the prothorax, invisible from above. The majority of these insects live in dry vegetable matter, such as in trunks and branches or under the bark of dead trees. Some *anobiìdeos* are common and very destructive pests. *Lasioderma serricorne* (Fabricius) is commonly known as the tobacco beetle, which attacks tobacco, museum specimens and collections of insects (Borror, 1969).

*Lasioderma serricorne* (F.) (Coleoptera: Anobiidae) is a well-known pest of a large number of stored products and the most serious pest of stored tobacco (Runner, 1919; Ashworth, 1993; Ryan, 1995; Levinson & Buchelos, 1988; Papadopoulou & Buchelos, 2002). According to Howe (1957), the development of *L. serricorne* depends on temperature and relative humidity of its environment and its diet. It is a common pest in heated buildings and multiplies in a wide variety of substrates. It infests soybeans, cocoa, chickpea, groundnut and coffee, among others (Alves, 2011).

The chickpea (*Cicer arietinum L*.) is a source of protein, carbohydrates, vitamins, fiber, minerals such as P, K, Ca, Mg, S, Na, Fe, Cu, Mn, Zn, and of 18 amino acids. (Ferreira et al., 2006).

## 2. MATERIALS AND METHODS

### 2.1. Material

The insects were obtained from the entomological biotherium of the Laboratory of Radiobiology and Environment of CENA-USP. They were fed with an experimental diet (control diet and chickpea diet), previously prepared and just when the first generation hatched completely, the experiments were conducted. This guarantees that the main cause of variation is the diet, since the doses of radiation and insect species were always the same. The randomly chosen control diet, consisted of three parts of wheat germ, one part of brewer’s yeast, and a slice of French bread toasted in an oven previously set up for humidity control. The diet of chickpeas consists only of whole grains crushed in a mechanical grinder to obtain flour.

### 2.2. Design of experiments

Only the descendants of the first generation from both new diets were used in experiments with radiation. The insects were raised on a diet of milled chickpeas made from only a mechanical grinder, or with the smallest possible degree of processing. The chickpea flour that emerged from this process was allocated in plastic containers with a cover of voal (soft tissue of polyester) for gas exchange in insects. The diets were radiosterilized prior to inoculation with a dose of 10 kGy to prevent mold and dust mites, other insects, and sometimes even the same insect that was already plaguing the food from the field. After inoculation, the insects developed in relative humidity (RH) and temperature environments for weeks under observation and care. In the emergence of a new generation, when there was enough population density, the experiments were conducted. The total number of insects in the experiment was 720 individuals split into 10 individuals per plate and three plates per treatment. The average temperature in the period was (21.4 ± 2.6) °C, the relative humidity in the same period was (62.0 ± 8.6) % and the averaged period of observation after irradiation was (37 ± 5) days. At emergence, the insects were placed in petri dishes in three replicates of ten adults per plate and three plates per dose, according to the proposed experimental design.

### 2.3 Irradiation Condition

The irradiations were conducted with ^60^Co gamma rays from a Gammacell source, model 220N of the Atomic Energy of Canada Limited, in the Technology Radiation Center of the IPEN, with doses of 5.0 Gy, 7.5 Gy, 10.0 Gy, 12.5 Gy, 15.0 Gy, 300.0 Gy, 600.0 Gy, 900.0 Gy, 1200.0 Gy and 1500.0 Gy, beyond their respective non-irradiated controls. The dose rate was 1.75 kGy/h and the source activity over the month was 2118.106 Ci. The sole biological parameter studied in this research was the survival of insects on the plates.

### 2.4 Experimental Model

The linear-quadratic model was the mathematical model adopted to fit the survival of insects as a function of the doses of ionizing radiation (Hall, 2000; Alpen, 2005; Kudryashov, 2008; Andrade & Bauermann, 2010). The statistical model of the computational analysis in SAS was the General Linear Model - GLM procedure in a completely randomized block design (Dean & Voss, 1999; Der & Everitt, 2002; Nogueira, 2007; Montgomery, 2005).

### 2.5 Statistical analysis

The analysis of variance was performed using the SAS version 9. The syntax of the SAS program for survival data analysis was developed based on Nogueira and Dean et al. Graphs were plotted in Origin software version 8.0.

## 3. RESULTS

The results showed different trends of the survival curves over time for the different diets, but for the same doses of radiation used. These differences can be visualized in graphs and are indicated in the analysis of variance. Table 1 shows the survival data of the control diet for doses up to 15 Gy, while table 2 presents the survival data for the control diet at doses from 300 to 1500 Gy.

**Table 1:** Survival values (mean ± SD) obtained for *Lasioderma serricorne*, fed with control diet and irradiated with doses of 5-15 Gy of ^60^Co.

| Day | Control | 5.0 Gy | 7.5 Gy | 10.0 Gy | 12.5 Gy | 15.0 Gy |
| --- | --- | --- | --- | --- | --- | --- |
| 0 | 10.0 ± 0.0 | 10.0 ± 0.0 | 10.0 ± 0.0 | 10.0 ± 0.0 | 10.0 ± 0.0 | 10.0 ± 0.0 |
| 4 | 9.7 ± 0.6 | 9.3 ± 1.2 | 9.0 ± 1.0 | 8.7 ± 1.2 | 9.7 ± 0.6 | 8.7 ± 1.2 |
| 8 | 7.0 ± 2.0 | 6.7 ± 1.2 | 7.0 ± 2.0 | 6.3 ± 2.1 | 8.0 ± 1.0 | 7.3 ± 2.3 |
| 12 | 5.7 ± 1.2 | 3.3 ± 1.2 | 6.0 ± 1.7 | 4.3 ± 1.2 | 7.0 ± 1.0 | 5.3 ± 2.3 |
| 16 | 3.7 ± 0.6 | 2.3 ± 0.6 | 3.0 ± 1.7 | 2.0 ± 1.0 | 3.3 ± 0.6 | 2.7 ± 2.1 |
| 20 | 1.3 ± 0.6 | 1.0 ± 1.0 | 1.3 ± 0.6 | 0.7 ± 1.2 | 2.3 ± 0.6 | 1.0 ± 1.7 |
| 24 | 0.7 ± 0.6 | 0.7 ± 1.2 | 0.7 ± 0.6 | 0.0 ± 0.0 | 2.0 ± 1.0 | 0.3 ± 0.6 |
| 28 | 0.7 ± 0.6 | 0.0 ± 0.0 | 0.7 ± 0.6 | 0.0 ± 0.0 | 1.3 ± 1.2 | 0.3 ± 0.6 |
| 32 | 0.0 ± 0.0 | 0.0 ± 0.0 | 0.7 ± 0.6 | 0.0 ± 0.0 | 0.3 ± 0.6 | 0.0 ± 0.0 |

**Table 2:** Values of survival (mean ± SD) obtained for *Lasioderma serricorne*, fed with control diet and irradiated with doses of 300-1500 Gy of ^60^Co.

| Day | Control | 300 Gy | 600 Gy | 900 Gy | 1200 Gy | 1500 Gy |
| --- | --- | --- | --- | --- | --- | --- |
| 0 | 10.0 ± 0.0 | 10.0 ± 0.0 | 10.0 ± 0.0 | 10.0 ± 0.0 | 10.0 ± 0.0 | 10.0 ± 0.0 |
| 4 | 9.7 ± 0.6 | 8.3 ± 0.6 | 7.7 ± 0.6 | 6.0 ± 2.0 | 6.7 ± 1.2 | 5.7 ± 3.0 |
| 8 | 7.0 ± 2.0 | 5.7 ± 2.1 | 3.7 ± 0.6 | 3.7 ± 3.0 | 3.3 ± 1.2 | 3.0 ± 2.6 |
| 12 | 5.7 ± 1.2 | 3.0 ± 1.7 | 1.7 ± 1.2 | 2.3 ± 2.1 | 2.0 ± 2.0 | 1.7 ± 1.2 |
| 16 | 3.7 ± 0.6 | 2.7 ± 1.5 | 1.0 ± 1.0 | 1.0 ± 1.0 | 1.7 ± 2.1 | 0.3 ± 0.6 |
| 20 | 1.3 ± 0.6 | 1.7 ± 0.6 | 0.3 ± 0.6 | 0.7 ± 0.6 | 0.3 ± 0.6 | 0.3 ± 0.6 |
| 24 | 0.7 ± 0.6 | 0.3 ± 0.6 | 0.3 ± 0.6 | 0.0 ± 0.0 | 0.0 ± 0.0 | 0.0 ± 0.0 |
| 28 | 0.7 ± 0.6 | 0.0 ± 0.0 | 0.0 ± 0.0 | 0.0 ± 0.0 | 0.0 ± 0.0 | 0.0 ± 0.0 |
| 32 | 0.0 ± 0.0 | 0.0 ± 0.0 | 0.0 ± 0.0 | 0.0 ± 0.0 | 0.0 ± 0.0 | 0.0 ± 0.0 |

Table 3 presents the survival data for the chickpea diet at doses from 5 to 15 Gy, while table 4 presents the survival data for the chickpea diet at doses from 300 to 1500 Gy.

**Table 3:** Values of survival (mean ± SD) obtained for *Lasioderma serricorne*, fed with chickpea diet and irradiated with doses of 5-15 Gy of ^60^Co.

| Day | Control | 5.0 Gy | 7.5 Gy | 10.0 Gy | 12.5 Gy | 15.0 Gy |
| --- | --- | --- | --- | --- | --- | --- |
| 0 | 10.0 ± 0.0 | 10.0 ± 0.0 | 10.0 ± 0.0 | 10.0 ± 0.0 | 10.0 ± 0.0 | 10.0 ± 0.0 |
| 4 | 10.0 ± 0.0 | 9.7 ± 0.6 | 10.0 ± 0.0 | 9.3 ± 1.2 | 9.0 ± 1.0 | 8.7 ± 0.6 |
| 8 | 10.0 ± 0.0 | 9.7 ± 0.6 | 9.7 ± 0.6 | 8.3 ± 0.6 | 7.7 ± 1.5 | 5.7 ± 0.6 |
| 12 | 8.3 ± 0.6 | 9.0 ± 1.0 | 9.7 ± 0.6 | 8.0 ± 1.0 | 6.0 ± 1.7 | 4.7 ± 0.6 |
| 16 | 5.7 ± 0.6 | 6.3 ± 3.0 | 6.7 ± 1.2 | 5.3 ± 2.1 | 3.7 ± 2.1 | 2.3 ± 0.6 |
| 20 | 2.3 ± 1.5 | 4.0 ± 1.7 | 4.0 ± 0.0 | 2.7 ± 2.1 | 1.3 ± 0.6 | 1.0 ± 1.0 |
| 24 | 0.7 ± 0.6 | 1.7 ± 0.6 | 1.7 ± 0.6 | 1.0 ± 1.0 | 1.0 ± 1.0 | 0.3 ± 0.6 |
| 28 | 0.3 ± 0.6 | 1.0 ± 1.0 | 0.3 ± 0.6 | 0.3 ± 0.6 | 0.3 ± 0.6 | 0.3 ± 0.6 |
| 32 | 0.0 ± 0.0 | 0.0 ± 0.0 | 0.3 ± 0.6 | 0.3 ± 0.6 | 0.0 ± 0.0 | 0.0 ± 0.0 |
| 36 | 0.0 ± 0.0 | 0.0 ± 0.0 | 0.3 ± 0.6 | 0.3 ± 0.6 | 0.0 ± 0.0 | 0.0 ± 0.0 |

**Table 4:**
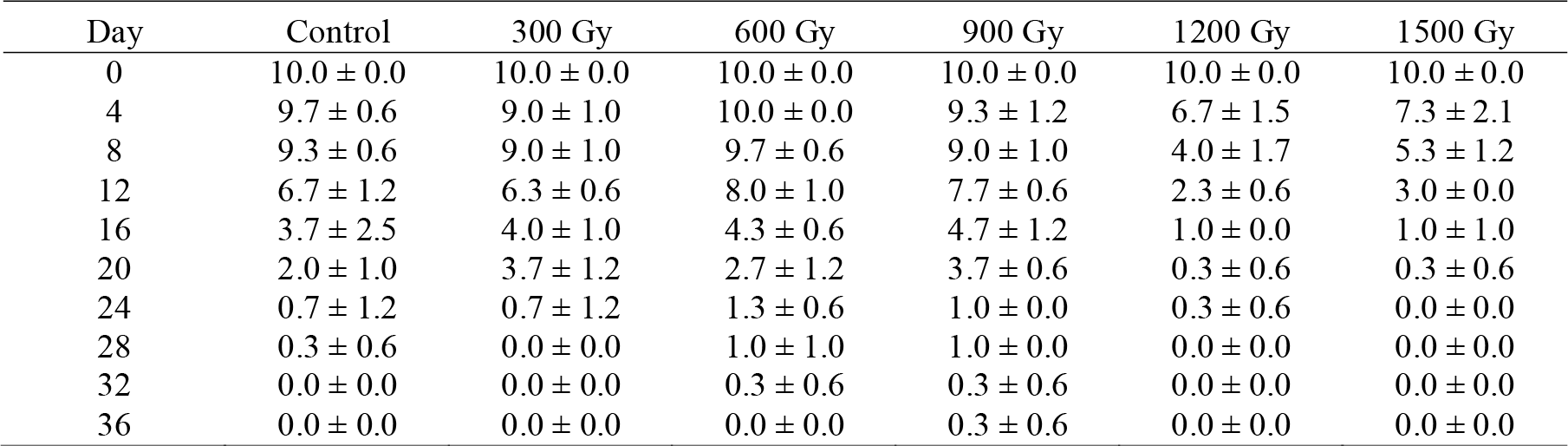
Values of survival (mean ± SD) obtained for Lasioderma serricorne, fed with chickpea diet and irradiated with doses of 300-1500 Gy of ^60^Co.

Figures 1 **and** 2 show the survival curve of the insects over time, fed with different diets and irradiated with several doses. For the lowest doses tested, it was not possible to verify significant differences between the control diet and chickpea diet in survival data up to 10 Gy, according to the Bonferroni’s test. From 12.5 Gy on, this difference appears only for the chickpea diet (p<0.0001).

**Figure 1:**
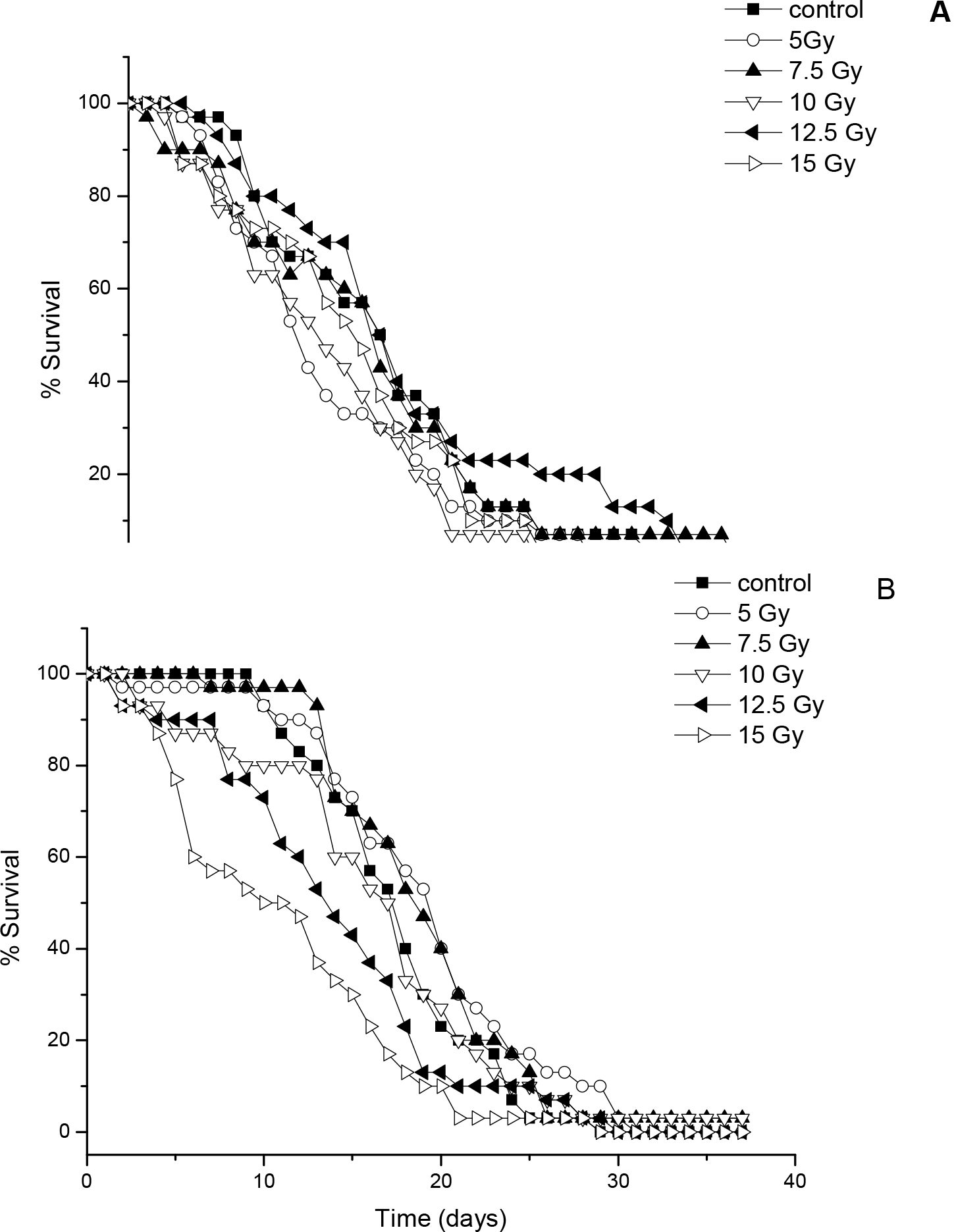
Survival curve of *Lasioderma serricorne* for doses in the range 5 - 15 Gy of ^60^Co. A) Insects fed with control diet. B) Insects fed with chickpea diet.

**Figure 2:**
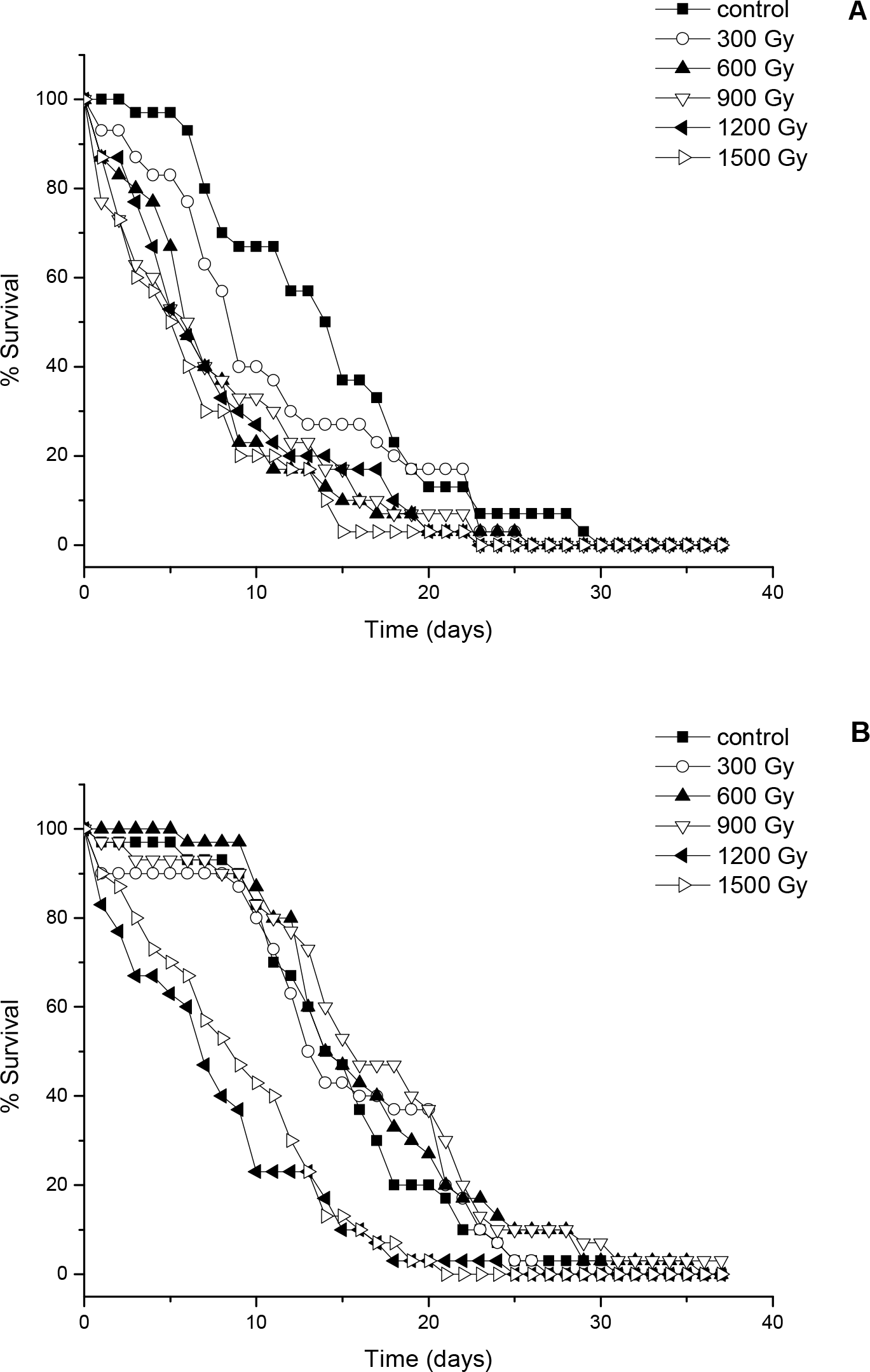
range 300 - 1500 Gy of ^60^Co: A) Insects in control diet. B) Insects in chickpea diet.

The results of the analysis of variance between the chickpea diet and control diet, for doses between 5 and 15 grays are shown in table 5.

**Table 5:** Analysis of variance for survival to doses in the range 5 - 15 grays between control diet and chickpea diet.

| Causes of Variation | DF | SS | MS | F<sub>calc</sub> | F<sub>crit</sub><br>(p =0.05) |
| --- | --- | --- | --- | --- | --- |
| Model | 45 | 5651.940 | 125.598 | 201.50 | 1.398 |
| Blocks | 11 | 265.634 | 24.149 | 38.74 | 1.807 |
| Treatments | 34 | 5386.307 | 158.421 | 254.15 | 1.468 |
| Error | 374 | 233.124 | 0.623 |  |  |
| Total | 419 | 5885.064 |  |  |  |

DF= degree of freedom ; SS= sum of squares ; MS=mean square ; F<sub>calc</sub> = calculated value of F ; F<sub>crit</sub> = tabulated value of F

The data obtained for insect survival were adjusted accordingly to the proposed mathematical model (R^2^ = 0.960) in table 5. The calculated value of F for the blocks is 38.74 and is much higher than the tabulated critical value of 1.807, indicating a highly significant difference due to a combination of diet and dose of radiation at the significance level of 95% (p<0.0001).

For 45 degrees of freedom of the model to 374 degrees of freedom of the residue (error), the F test indicated a highly significant result (F_calc_ = 201.50 ≫ F_crit_ = 1.398; p<0.0001). The result of the analysis of variance for the influence due to treatments (time – independent variable, F_calc_ = 254.15 ≫ F_crit_ = 1.468; p<0.0001) and the influence on the results due to blocks (combination of diet and radiation dose – dependent variable, F_calc_ = 38.74 > F_crit_ = 1.807; p<0.0001) are both (time and diet) significant according to data organized in table 4 for a level of significance of 95%.

The results of analysis of variance between chickpea diet and control diet to doses in the range of 300 to 1500 Gy are organized in table 6.

**Table 6:**
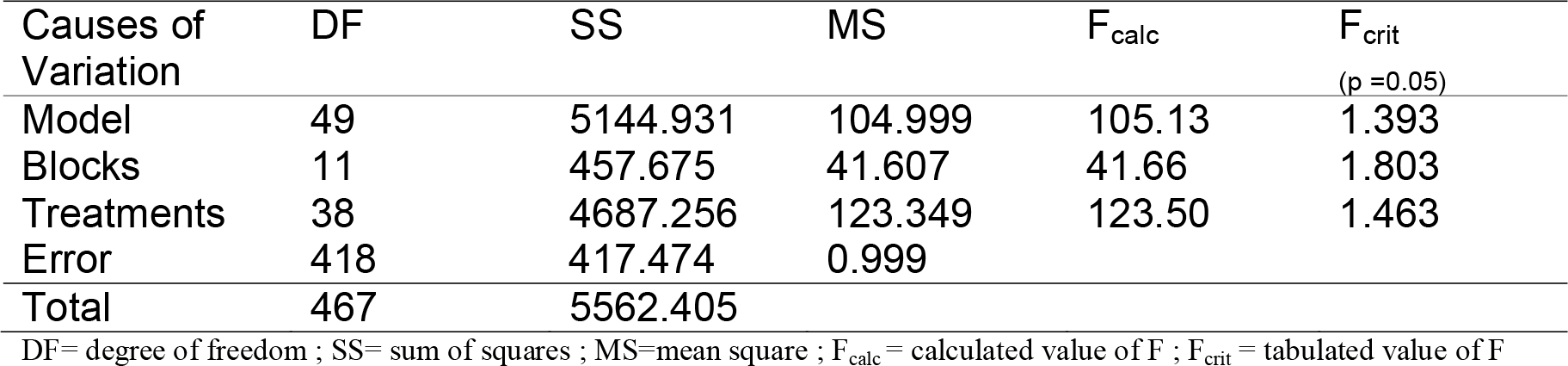
Analysis of variance for survival to doses in the range 300 - 1500 grays between control diet and chickpea diet.

The coefficient of determination of the model in table 6 is R^2^ = 0.925, which indicates an excellent correspondence between the data and the proposed model of analysis. The calculated value of F for the blocks is 41.66, much larger than the tabulated critical value which is 1.807, indicating a highly significant difference due to a combination of both diet and radiation dose at the significance level of 95% (p<0.0001).

For 49 degrees of freedom of the model to 418 degrees of freedom of the residue (error), the F test indicated a highly significant difference (F_calc_ = 105.13 ≫ F_crit_ = 1.393; p<0.0001). The result of analysis of variance for the influence due to treatments (time – independent variable, F_calc_ = 123.50 ≫ F_crit_ = 1.463; p<0.0001) and the influence on the results due to blocks (combination of diet and radiation dose – dependent variable, F_calc_ = 41.66 > F_crit_ = 1.803; p<0.0001) are both significant according to data organized in table 4 for a level of significance of 95%.

## 4. DISCUSSION

This research represents a preliminary study for the development and enhancement of a technical investigation on the possible presence of natural radioprotective agent in foods. Some parameters can be improved and optimized, but the methodology has demonstrated to be functional. There is the possibility that nutritional factors cause changes in the survival response of a population, however the doses tested are relatively high (300 to 900 Gy), which may indicate another possibility, namely the radioprotective agent action. We would like to propose here a new research line in general radiation biology that can be called “natural radioprotectors in foods.” We could observe in other experiments in progress in this laboratory not yet published, that other foods may be tested using this methodology. Brazil is one of the most promising countries in the world for the development of these research proposals due to its rich natural dietary diversity.

To infer possible predictions of the results obtained in this experiment with insects to the mammalian class, it is important to remember that the taxonomic group of insects is generally more radioresistant than mammalians (Linsley, 1997), wich may mean that mammalian cells could present a similar behavior for much smaller doses. The importance of this study is the new approach technique developed to investigate the presence of radioprotectors in food and the result obtained with the radioprotective effects of chickpea, which are not reported in the relevant literature.

This is not the first work of research showing evidences of natural radioprotectors in food, but it is the first to focus on chickpeas. A-bomb survivors who frequently consumed “miso” (Japanese soybean fermented paste) demonstrated decreased radiation damage. Because of this report, Europeans were recommended to eat miso after the Chernobyl accident (Ohara, 2001). Curcumin, a culinary spice from Vedic culture in India, has been found to exert a dual mode of action, as radioprotector or radiosensitizer after irradiation, depending on its concentration (Jagetia, 2007a). Spigoti et al. (2011) investigated the radioprotective effects of Brazilian propolis AF 08 in Chinese hamster ovary (CHO-K1) and prostate tumor cells (PC 3) irradiated with ^60^Co. Analyzing micronucleus induction, cellular viability and clonogenic death, they verified the radioprotective and non-toxic effect of propolis at concentrations of 5 - 100 µg/ml for a dose range of 1 - 4 Gy. Jagetia (2007b) has studied the radioprotective effect of several plants such as *Gingko biloba*, *Panax ginseng*, *Piper longum*, and *Zingiber officinale*, among others and found that they protect against radiation-induced lethality, lipid peroxidation, and DNA damage. For doses of 10, 12.5 and 15 Gy, we can see a progressive distance from control curve, as would be expected. In figure 2B, the graph shows a clear division between two groups. The first graph includes the control. At doses of 300, 600 and 900 Gy, which are relatively high, the curves move farther from the control in a path approximately proportional to the dose, but this did not occur for very high doses of 1200 Gy and 1500 Gy. In the control diet (figure 2A) this expected separation of the curves is noticeable.

Applying the Bonferroni test to the survival data from insects reared on control diet, we could observe a significant difference between 300, 600, 900 Gy and the control, as expected. However for the chickpea diet, there was no significant difference between 300, 600, 900 Gy and the control, even though there should have been. This can be seen in figure 2. Applying the Bonferroni test for high doses, however, there was no significant difference between 1200 and 1500 Gy in both diets. There is a significant difference between the survival number of insects for these doses compared to their controls for both diets. This may indicate that for very high doses, the possible radioprotective effect fails.

This situation can be explained by the presence of a radioprotective agent. We suggest a further study of the nutritional composition of chickpeas with HPLC to investigate the possible presence of already known radioprotective agents. We also suggest that the same experiment be developed for high-LET radiation as a neutron beam in a nuclear research reactor.

## 5. CONCLUSIONS

We presented statistical evidence that the chickpea diet has radioprotective properties in the insect for gamma rays. This research methodology also showed some important advantages over the experiments with mice: low cost, small space requirement, the possibility of using a large number of individuals as statistical security and easy counting of insect individuals. The main problem found was the infestation of some diets by mites, which are common in tropical countries, these forced some repetitions of the trials.

## 6. ACKNOWLEDGEMENTS

To the National Commission of Nuclear Energy (CNEN) for providing facilities during the development of this research; To researcher João Justi from the Biological Institute of São Paulo for the initial suggestions about preparing the diets; To the CENA-USP for providing the insect matrices; To IPEN which offered all the infrastructure for laboratory studies with mammalian cells and made adaptations to undertake the work with insects; To technicians Carlos Silveira and Elizabeth Somessari of the CTR for their special care in insect irradiation; To Laura Li from Fulbright Scholar / U.S.-Brazil Fulbright Commission for reviewing English grammar; To researcher Daniel Perez for the suggestions and comments; and to Ottília Benzi (*In memorian*) for their support and motivation.

